# Luteal-phase resistance training enhances nocturnal heat dissipation and delta power during sleep

**DOI:** 10.1101/2025.10.21.676688

**Authors:** Momo Fushimi, Ryusei Iijima, Shiori Noguchi, Aoi Kawamura, Kenichi Kuriyama, Masako Tamaki, Sayaka Aritake-Okada

## Abstract

**Study Objectives:** This study examined the effects of resistance training on sleep architecture, physiological heat dissipation, and δ power in young adult women during the follicular and luteal phases of the menstrual cycle.

**Methods:** Nine healthy young women participated in a four-condition crossover protocol comprising (1) follicular phase non-exercise, (2) follicular phase exercise, (3) luteal phase non-exercise, and (4) luteal phase exercise. The exercise condition consisted of 30 minutes of resistance training at 70% of one-repetition maximum performed during the day. During each night, electroencephalography, body temperature, and core body temperature were measured in the home environment. The distal–proximal body temperature gradient (DPG), which is a validated indicator of heat dissipation, was calculated

**Results:** Resistance training enhanced heat dissipation, as reflected by increased DPG values, and increased the proportion of stage N3 sleep during both menstrual cycle phases, with more pronounced effects during the luteal phase. Sleep from bedtime to wake time was divided into four equal segments. Under the luteal phase with exercise, the appearance of stage N3, δ power, and the DPG were elevated in the mid-to-late segments of sleep.

**Conclusions:** These findings suggest that daytime resistance training promotes nocturnal deep sleep and facilitates thermoregulatory heat loss, as indicated by an increased DPG, particularly during the luteal phase when thermoregulation is less stable than the follicular phase. This training may represent a practical intervention to improve sleep quality and physiological recovery in women across different phases of the menstrual cycle.

**STATEMENT OF SIGNIFICANCE:** Women often experience sleep disturbances during the luteal phase of the menstrual cycle, but the physiological mechanisms and effective interventions remain unclear. This study shows that daytime resistance training enhances nighttime heat dissipation and increases slow-wave sleep during this phase. These findings suggest that exercise is a practical approach to alleviate menstrual cycle-related sleep issues. By clarifying how exercise interacts with thermoregulation and sleep across the cycle, this work contributes to understanding female-specific sleep physiology and supports the development of non-pharmacological strategies to improve women’s sleep health in women. Future research should examine the long-term effects of daytime resistance training and compare different exercise modalities across diverse populations.

## INTRODUCTION

Blood concentrations of female hormones increase or decrease with each menstrual cycle phase[1]. The core body temperature increases by 0.3℃–0.7℃ in the luteal phase, when progesterone concentrations are higher than those in the follicular phase before ovulation[2]. Body temperature rhythms associated with the menstrual cycle phase are closely related to sleep[3]. Heat dissipation from the periphery and a decrease in core body temperature are essential for smooth sleep onset and maintenance. However, during the luteal phase, not only is core body temperature elevated, but the amplitude of the core body temperature rhythm is also diminished, potentially disrupting the balance between wakefulness and sleep[2].

Elevated progesterone concentrations in the luteal phase have been reported to cause strong daytime sleepiness [3,4]. An elevation in progesterone concentrations also reduces sleep quality by suppressing the heat dissipation response at sleep onset during the night, causing difficulty in sufficiently lowering core body temperature, increasing wake after sleep onset [5], and reducing REM(rapid eye movement) sleep and slow-wave sleep [6]. If there is a considerable degree of sleep disturbance in the luteal phase, it may be one of the symptoms of premenstrual syndrome (PMS), a premenstrual dysphoric mood disorder, which can lead to serious problems in a woman’s social life [7–9].

Exercise is a widely recognized non-pharmacological intervention for improving sleep quality. Previous studies have shown that various types of physical activity, including resistance and aerobic training, can enhance objective sleep parameters[10,11]. In women, daytime exercise enhances heat dissipation responses during wakefulness, particularly during the luteal phase [12]. Moreover, habitual exercisers have reportedly shown less heat loss suppression during the mid-luteal phase[13]. Exercise has also been shown to alleviate subjective symptoms of sleeplessness in women with PMS [14].

Our previous study showed that daytime physical activity promoted heat dissipation and improved slow-wave sleep in healthy young men[11]. Similar effects may occur in women, although hormonal fluctuations across the menstrual cycle phase may modulate the response to exercise. In particular, the effect of exercise on nocturnal thermoregulation and sleep architecture may differ between the follicular and luteal phases. Despite these possibilities, few studies have examined how exercise-induced heat dissipation affects sleep in women across stages of the menstrual cycle.

We hypothesized that moderate intensity resistance training enhances pre-sleep and nocturnal heat dissipation, particularly in the luteal phase, when thermoregulatory function is impaired. Furthermore, based on our original protocol, we hypothesized that this enhancement of heat dissipation would promote slow-wave sleep. This study aimed to investigate the effects and physiological mechanisms of an approximately 30-minute moderate intensity resistance training session on heat dissipation and objective and subjective sleep parameters in healthy young women during the follicular and luteal phases of the menstrual cycle. Clarifying these mechanisms could contribute to our understanding of menstrual-phase-specific thermoregulatory physiology and inform strategies to alleviate menstrual cycle phase-related sleep disturbances.

## METHODS

### Participants

This study was approved by the Ethical Review Committee of Saitama Prefectural University (Approval No. 19024). All participants provided written informed consent after receiving a full explanation of the study’s aims, procedures, and potential risks. Twelve healthy young adult female university students (mean age: 20.83 ± 0.94 years) were recruited via posters and university email lists. The inclusion criteria were a regular menstrual cycle, habitual sleep–wake schedules (bedtime between 23:00 and 01:00 hours), and no habitual physical exercise. The exclusion criteria included the following: diagnosed sleep, psychiatric, or neurological disorders; shift work; recent travel across time zones (within 1 month); and current use of medications or substances (e.g., caffeine, nicotine, and/or alcohol) that could affect sleep or body temperature.

Sleep–wake patterns were verified with actigraphy (HJA-750C; Omron, Kyoto, Japan) for 1 month before the experiment. The participants completed the following screening questionnaires: Pittsburgh Sleep Quality Index[15], General Health Questionnaire[16], The Center for Epidemiologic Studies Depression Scale[17], and Menstrual Distress Questionnaire[18].

### Experimental protocol

The participants completed the four following experimental sessions in a randomized, crossover design: (1) follicular phase non-exercise, (2) follicular phase exercise, (3) luteal phase non-exercise, and (4) luteal phase exercise sessions. The order of menstrual cycle phases (follicular or luteal) was randomized, while within each phase, the non-exercise session always preceded the exercise session. Follicular phase sessions were scheduled 7–10 days after the onset of menstruation, and luteal phase sessions were scheduled 19–22 days after[12]. Each session was spaced at least 2 weeks apart. To confirm the menstrual cycle phase, daily basal body temperature recordings were collected for at least 1 month before the experiment. The menstrual cycle phase was tracked for 1 month via daily basal body temperature recordings (CTEB503L-E; Citizen, Tokyo, Japan). When necessary, ovulation prediction kits (Do Test LH Ovulation Prediction Test Stick Type; Rohto Pharmaceutical, Osaka, Japan) were used to confirm phase timing. On the day of testing, the participants refrained from physical activity (confirmed via actigraphy and self-reporting) and arrived at the laboratory at 13:00 hours. In control conditions, the participants read books quietly for 30 minutes beginning at 14:00 hours, and in exercise conditions, they completed a bodyweight resistance training protocol session (Figure 1). The session included six resistance training exercises (push-ups, sit-ups, hip lifts, lunges, back extensions, and half squats), which were performed at 70% of one-repetition maximum [19], with 2-minute rest intervals, guided by a metronome (96 BPM(beats per minuite)). The individual one-repetition maximum was determined by a certified athletic trainer at least 1 week previously. The participants returned home at 17:00 hours. The participants wore actigraphy devices to verify wakefulness until bedtime.

**Figure 1.**
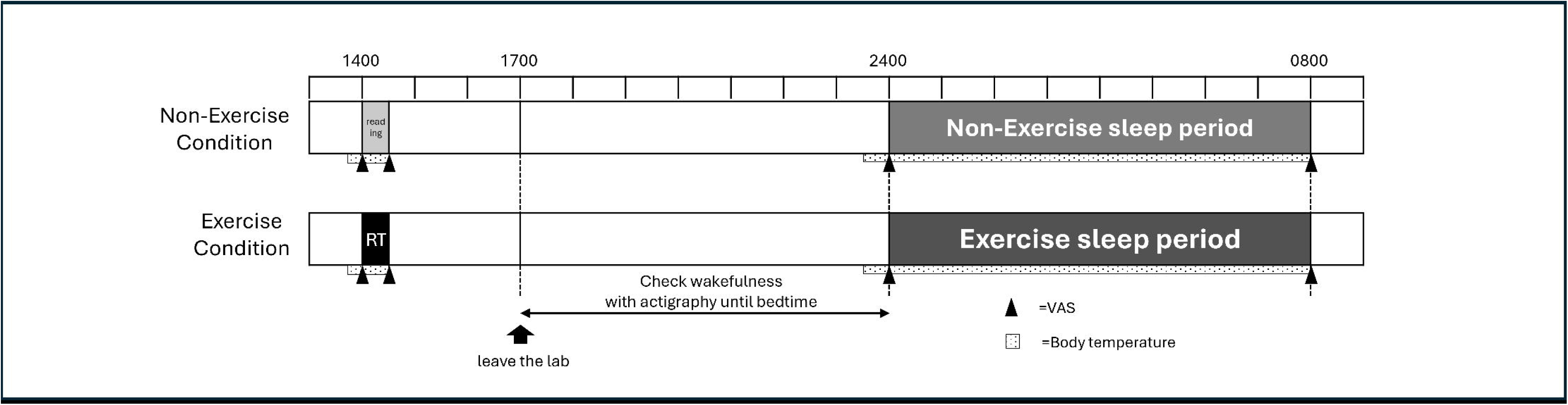
Experimental schedule and protocol overview. Participants arrived at the laboratory at 13:00 and engaged in either 30 minutes of reading (non-exercise condition) or resistance training (RT) at moderate intensity (exercise condition) beginning at 14:00. After completing the session, they left the laboratory at 17:00 and returned home. Wakefulness until bedtime was monitored using an actigraphy device. Nocturnal sleep was recorded for approximately 8 hours at home, starting at each participant’s habitual bedtime. Distal and proximal skin temperatures and estimated core body temperature were continuously recorded at 1-minute intervals throughout the experimental day, and are illustrated as a bar with a dotted pattern in the figure. Subjective sleep quality was assessed using a Visual Analogue Scale (VAS) at four time points: before and after reading or RT, at bedtime, and upon awakening (▴). A randomized crossover design was used to counterbalance the order of menstrual phase (follicular and luteal) and condition (non-exercise and exercise) across participants. When both conditions had to be completed within the same menstrual phase due to scheduling constraints, the non-exercise condition was conducted first to prevent carryover effects of RT on subsequent sleep measurements.

### Sleep and temperature measurements

Overnight sleep of the participants was recorded during their habitual sleep period at home using a portable sleep electroencephalography (EEG) system (ZA-IX and ZA-X; Proassist, Osaka, Japan) with Fp1–A2 electrode placement, accompanied by electrooculography and surface electromyography of the chin. The high-cut and low-cut filter settings were set to 35 Hz and 0.3 Hz for EEG and electrooculography, and to 100 Hz and 10 Hz for electromyography, respectively. Sleep stages were initially scored by the polysomnographic technologists and then subsequently reviewed and finalized by a registered polysomnographic technologist, certified by the Board of Registered Polysomnographic Technologists in the U.S., in 30-second epochs in accordance with the American Academy of Sleep Medicine Scoring Manual Version 2.5[20]. The following parameters were calculated: time in bed, sleep period time, total sleep time, wake after sleep onset, sleep efficiency, sleep onset latency, durations and percentages of each sleep stage (N1, N2, N3, and REM), and stage latencies. To analyze temporal dynamics of sleep architecture, each subject’s sleep period from sleep onset to wake time was divided into four equal segments, and the percentage of each sleep stage was calculated for each segment.

A continuous spectral analysis was performed to examine the temporal dynamics of δ (0.5–4.0 Hz) EEG power during nocturnal sleep. This analysis was conducted using continuous spectral analysis Play Analysis software (NoruPro, Tokyo, Japan), which applies a fast Fourier transform-based algorithm to consecutive 30-second epochs. In each subject, the cumulative total δ power was calculated for the entire sleep period (sleep onset to wake time) and for each of four equal segments, which were obtained by dividing the total sleep period into quartiles. Values are expressed in μV², enabling assessment of overall and segment-specific slow-wave activity.

Body temperatures were recorded every minute at distal (the back of the hand and the dorsum of the foot) and proximal (forehead and subclavian) sites using button-type sensors (Thermocron SL; KN Laboratories, Osaka, Japan). The tympanic membrane temperature[2] was also recorded every minute (LT-8; Gram Corporation, Saitama, Japan). The distal–proximal body temperature gradient (DPG), which is a validated index of heat dissipation[21], was calculated as the distal minus proximal temperatures. The DPG was computed every 2 minutes during the 20 minutes before bedtime, and every 15 minutes during sleep.

Subjective sleep-related variables, including sleep latency, sleepiness, fatigue, and restorative sleep, were measured using a visual analog scale and the Stanford Sleepiness Scale [22] at four time points: pre– and post-intervention, bedtime, and upon waking.

### Statistical analysis

The Wilcoxon signed-rank test was used to compare subjective and objective sleep parameters, body temperature indices, and DPG values between conditions. Spearman’s rank correlation was used to examine associations between the DPG and sleep parameters. Statistical analyses were performed using SPSS version 28.0 (IBM, Armonk, NY, USA). Results are reported as the mean ± standard deviation or standard error, with statistical significance set at *p* < 0.05.

## RESULTS

### Participant characteristics

Thirty-six data from nine participants (mean age: 21.00 ± 1.07 years) were obtained in the present study. Three participants were excluded because of (1) EEG communication failure, (2) loss of body temperature data from a tape-induced rash, and (3) an incorrect self-reported bedtime. The participants’ characteristics are shown in Table 1. The basal body temperature was significantly higher in the luteal phase than in the follicular phase (*p* = 0.028). Menstrual symptoms were assessed using the Menstrual Distress Questionnaire, also showed significantly higher scores in the luteal phase, indicating a typical cyclic pattern (*p* < 0.05). However, the symptom severity remained within a mild range. No significant phase difference was observed for daytime sleepiness assessed by the Stanford Sleepiness Scale (*p* = 0.071)).

**Table 1.**
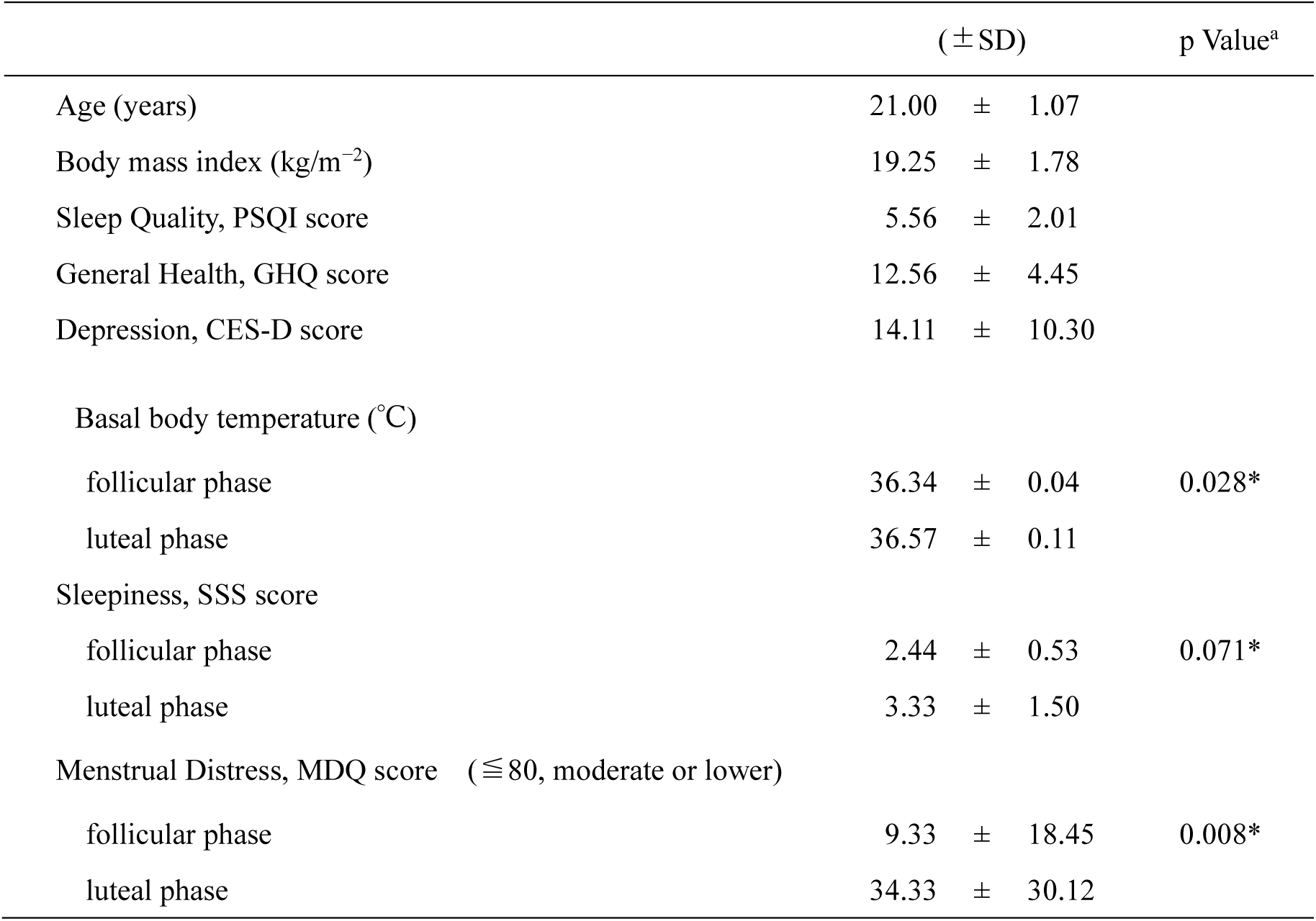
Participant characteristics and menstrual phase–related variables. Descriptive statistics for demographic, psychological, and physiological measures, including age, BMI, PSQI, GHQ, CES-D, basal body temperature, SSS, and MDQ scores during the follicular and luteal phases. a Values for basal body temperature, SSS, and MDQ represent comparisons between the follicular phase baseline (FPbase) and luteal phase baseline (LPbase) conditions (*p <* 0.05).

### Objective sleep parameters

Table 2 shows the objective sleep parameters during nocturnal sleep. In the follicular phase, the exercise condition was associated with a significantly lower percentage of stage N1 sleep (z = 2.033, *p* = 0.042), a higher percentage of stage N3 sleep (z = 2.240, *p* = 0.025), and a shorter latency to stage N3 sleep (z = 2.240, *p* = 0.025) than in the non-exercise condition. In the luteal phase, the exercise condition resulted in a significantly shorter duration of stage N2 (z = 2.197, *p* = 0.028), longer duration of stage N3 (z = 2.201, *p* = 0.028), lower percentage of stage N2 (z = 2.366, *p* = 0.018), and higher percentage of stage N3 (z = 2.197, *p* = 0.028) sleep than in the non-exercise condition. No significant differences were observed in other objective parameters between the two conditions.

**Table 2.** Objective sleep parameters under non-exercise and exercise conditions during the follicular and luteal phases. Mean values (± SD) of objective sleep measures across non-exercise and exercise conditions during the follicular and luteal phases. Parameters include sleep timing, duration, efficiency, sleep stage percentage, and latency. a Comparison between follicular-phase non-exercise and exercise conditions (*p <* 0.05). b Comparison between luteal-phase non-exercise and exercise conditions (*p <* 0.05).

| Objective sleep parameters | FP non-exercise<br>condition (± SD) | FP exercise<br>condition (± SD) | p Value<sup>a</sup> | LP non-exercise<br>condition(± SD) | LP exercise<br>condition (± SD) | p Value<sup>b</sup> |
| --- | --- | --- | --- | --- | --- | --- |
| bedtime | 24:49:39 ± 0:30:10 | 24:53:19 ± 0:33:31 | 0.833 | 24:37:36 ± 0:19:56 | 24:30:57 ± 0:23:46 | 0.600 |
| time in bed (min) | 416.25 ± 73.08 | 407.33 ± 90.17 | 0.779 | 442.86 ± 69.37 | 387.49 ± 106.20 | 0.063 |
| sleep period time (min) | 407.55 ± 74.12 | 393.10 ± 102.42 | 0.484 | 417.74 ± 62.32 | 379.29 ± 109.67 | 0.128 |
| total sleep time (min) | 383.93 ± 72.07 | 382.85 ± 98.28 | 0.674 | 386.53 ± 71.18 | 366.93 ± 112.87 | 0.398 |
| sleep efficiency (%) | 92.31 ± 6.65 | 93.38 ± 7.19 | 0.575 | 88.03 ± 13.24 | 93.77 ± 4.85 | 0.128 |
| sleep latency (min) | 9.19 ± 6.92 | 8.20 ± 6.22 | 0.674 | 9.68 ± 6.29 | 6.84 ± 5.61 | 0.345 |
| WASO (min) | 24.89 ± 34.61 | 18.09 ± 20.55 | 0.672 | 33.64 ± 50.75 | 15.14 ± 7.59 | 0.398 |
| stageN1 duration (min) | 23.40 ± 9.87 | 18.75 ± 9.01 | 0.161 | 24.86 ± 8.00 | 20.93 ± 4.98 | 0.176 |
| stageN2 duration(min) | 199.00 ± 55.68 | 185.98 ± 63.48 | 0.327 | 235.17 ± 62.08 | 162.24 ± 68.99 | 0.028* |
| stageN3 duration (min) | 90.00 ± 21.93 | 112.50 ± 20.49 | 0.050 | 75.43 ± 19.77 | 107.93 ± 30.95 | 0.028* |
| stageR duration (min) | 71.53 ± 33.68 | 65.63 ± 42.08 | 0.674 | 51.07 ± 29.12 | 75.83 ± 62.02 | 0.237 |
| stageN1/TST (%) | 6.14 ± 2.29 | 4.86 ± 1.64 | 0.042* | 6.44 ± 1.88 | 6.00 ± 1.49 | 0.397 |
| stageN2/TST (%) | 51.66 ± 9.95 | 48.29 ± 10.27 | 0.484 | 60.51 ± 10.00 | 43.87 ± 9.57 | 0.018* |
| stageN3/TST (%) | 24.14 ± 7.20 | 30.96 ± 9.31 | 0.025* | 19.59 ± 4.36 | 32.01 ± 12.79 | 0.028* |
| stageR/TST (%) | 18.06 ± 8.12 | 15.91 ± 9.17 | 0.484 | 13.43 ± 8.30 | 18.14 ± 13.68 | 0.128 |
| stageN1 latency (min) | 8.88 ± 7.92 | 6.39 ± 6.08 | 0.401 | 8.77 ± 6.29 | 5.41 ± 5.43 | 0.173 |
| stageN2 latency (min) | 10.06 ± 7.79 | 9.76 ± 6.92 | 0.208 | 27.97 ± 45.36 | 7.70 ± 5.64 | 0.204 |
| stageN3 latency (min) | 25.94 ± 7.58 | 19.83 ± 9.16 | 0.025* | 45.76 ± 45.28 | 17.91 ± 7.23 | 0.063 |
| stageR latency (min) | 131.69 ± 42.01 | 107.54 ± 33.47 | 0.499 | 126.76 ± 48.54 | 88.63 ± 30.55 | 0.128 |

### Body temperature, DPG, and tympanic temperature

Figure 2 shows changes in the time course of body temperature, the DPG, and tympanic temperature before and during bedtime. In the luteal phase, the DPG was significantly higher in the exercise condition than in the non-exercise condition at −6, −4, −2, and 0 minutes before bedtime (z = 1.965, *p* = 0.049; z = 2.366, *p* = 0.018; z = 2.120, *p* = 0.034; and z = 2.371, *p* = 0.018, respectively). A pointwise analysis showed a significantly elevated DPG at 4.5 hours after bedtime in the exercise condition compared with the non-exercise condition (z = 2.028, *p* = 0.043). In contrast, during the follicular phase, no significant difference in the DPG was observed between the exercise and non-exercise conditions at any time point. In addition, no significant difference in tympanic temperature was found between conditions before or after bedtime in either menstrual cycle phase.

**Figure 2.**
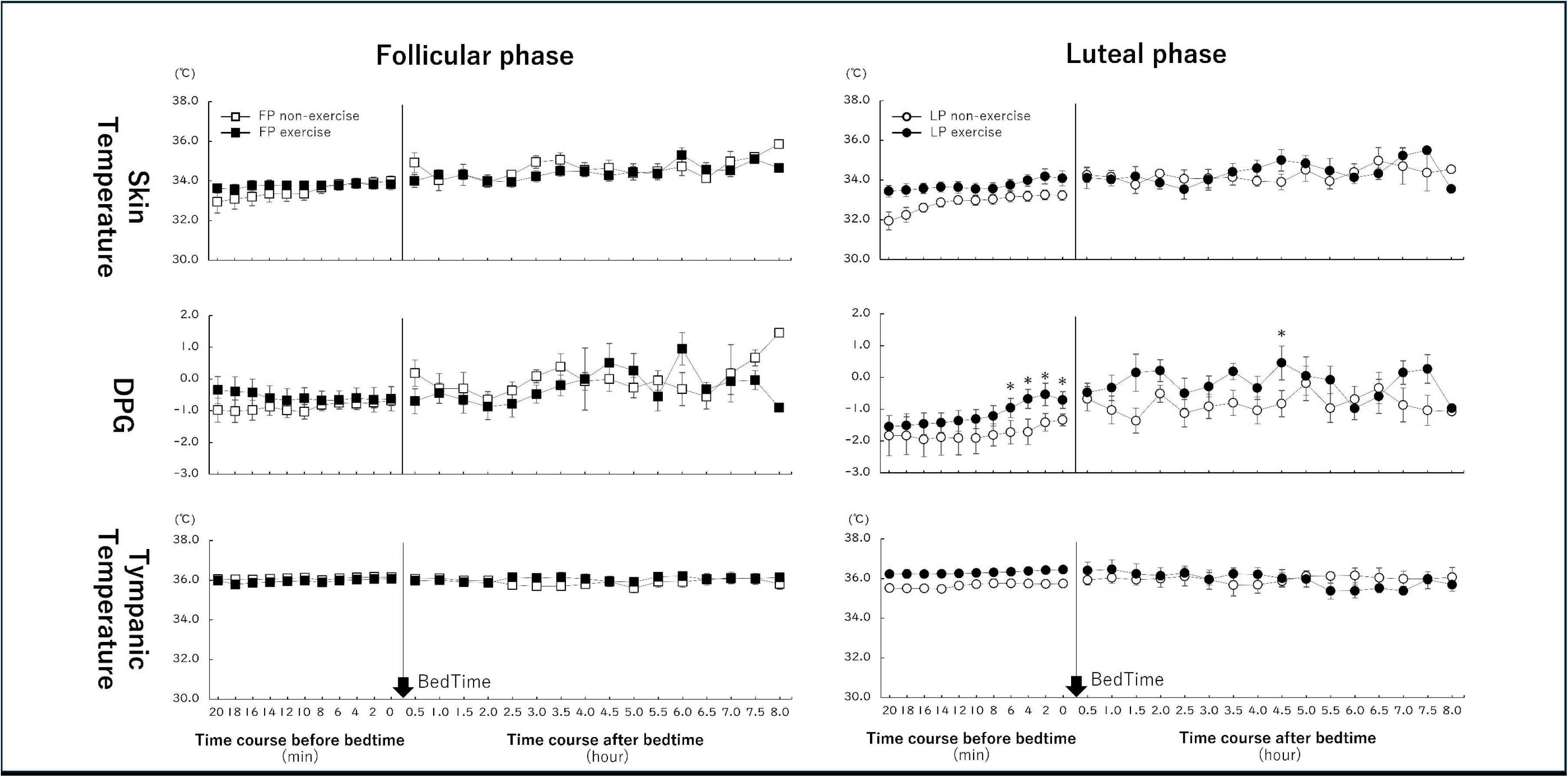
Time course of thermoregulatory responses before and after bedtime in each menstrual phase. Time series data of distal skin temperature (top panels), distal–proximal skin temperature gradient (DPG; middle panels), and tympanic temperature (bottom panels) measured before and after habitual bedtime during the follicular (left) and luteal (right) phases. Open symbols represent the non-exercise condition (□: FP, ○: LP), and filled symbols represent the exercise condition (▪: FP, ●: LP). Bedtime (vertical line) marks the transition from the pre-sleep period (minutes) to the post-sleep period (hours). Values are presented as mean ± SE. Asterisks indicate significant differences between exercise and non-exercise conditions at each time point (*p <* 0.05).

### Temporal distribution of sleep stages (four-segment analysis)

Each subject’s sleep period (from sleep onset to wake time) was divided into four equal segments, and the percentage of each sleep stage within each segment was calculated (Figure 3). The total duration of each sleep stage across the entire night is shown in Figure 3 as a visual supplement to Table 2, which presents detailed numerical values. In the luteal phase, the percentage of stage N2 sleep was significantly lower and that of stage N3 sleep was significantly higher in the third segment in the exercise condition compared with the non-exercise condition (z = 2.366, *p* = 0.018 and z = 2.028, *p* = 0.043, respectively). In the follicular phase, the percentage of stage W sleep in the first segment was significantly lower in the exercise condition than in the non-exercise condition (z = 2.366, *p* = 0.018).

**Figure 3.**
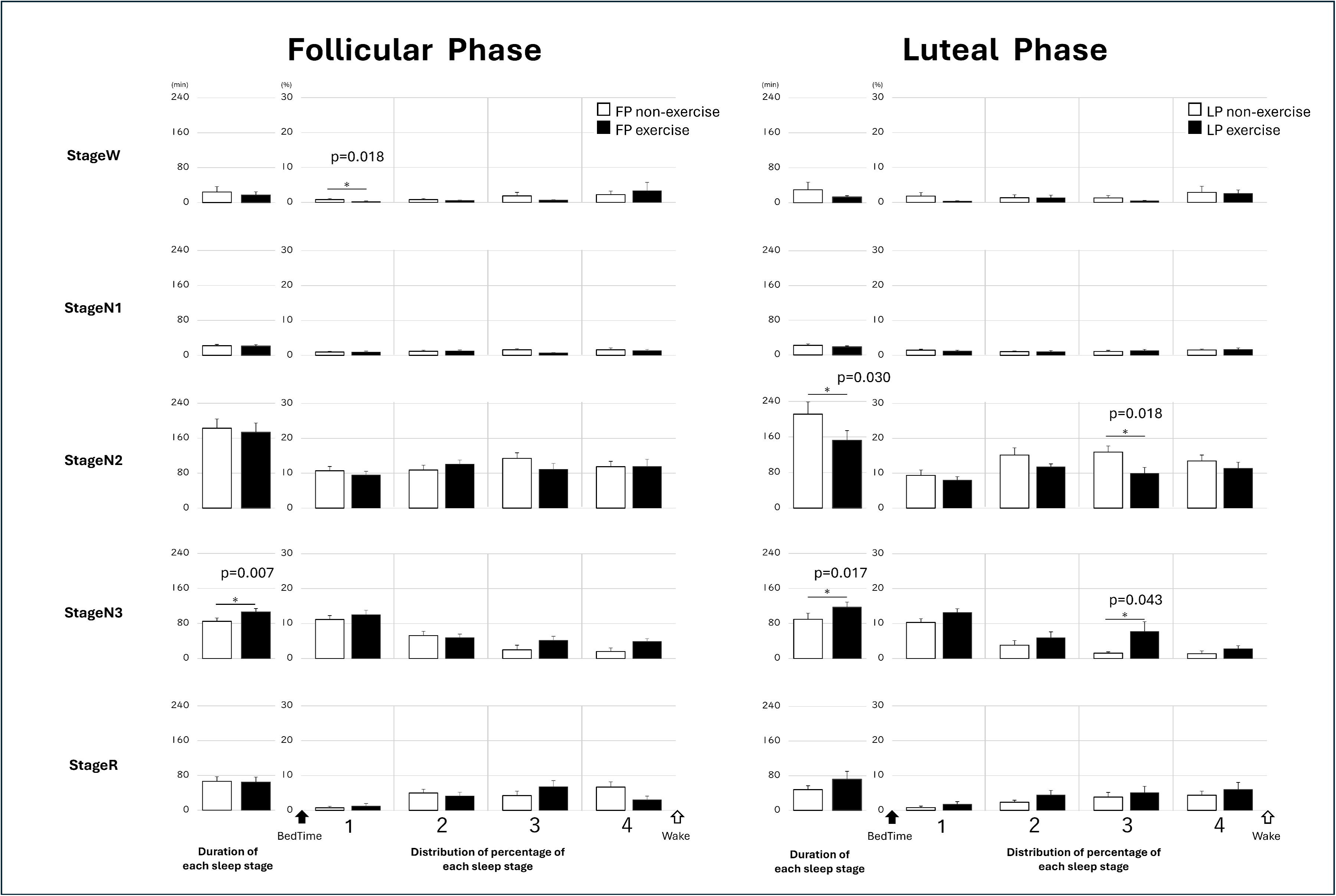
Effects of exercise on sleep stage distribution and timing in the follicular and luteal phases. Relative percentage (right four bars) of each sleep stage—W, N1, N2, N3, and R—during nocturnal sleep in the follicular (left panels) and luteal (right panels) phases under non-exercise (open bars) and exercise (filled bars) conditions. The distribution of each sleep stage was analyzed across four equal quartiles of total sleep time, from bedtime (1) to wake (4). Values represent mean ± SE. Asterisks and p-values indicate significant differences between conditions (*p <* 0.05).

### δ Power and DPG distribution during sleep

We examined the distribution of δ power (0.5–4.0 Hz) and the DPG during stage N3 sleep because of the increased occurrence of stage N3 sleep in the latter half of nocturnal sleep during the luteal phase exercise condition (Figure 4). In the luteal phase, δ power and the DPG were significantly higher in the third segment in the exercise condition than in the non-exercise condition (z = 2.028, *p* = 0.043 and z = 1.922, *p* = 0.046, respectively). In the follicular phase, δ power was significantly higher in the fourth segment during the exercise condition than in the non-exercise condition (z = 2.380, *p* = 0.017), while the DPG was not significantly different between the conditions.

**Figure 4.**
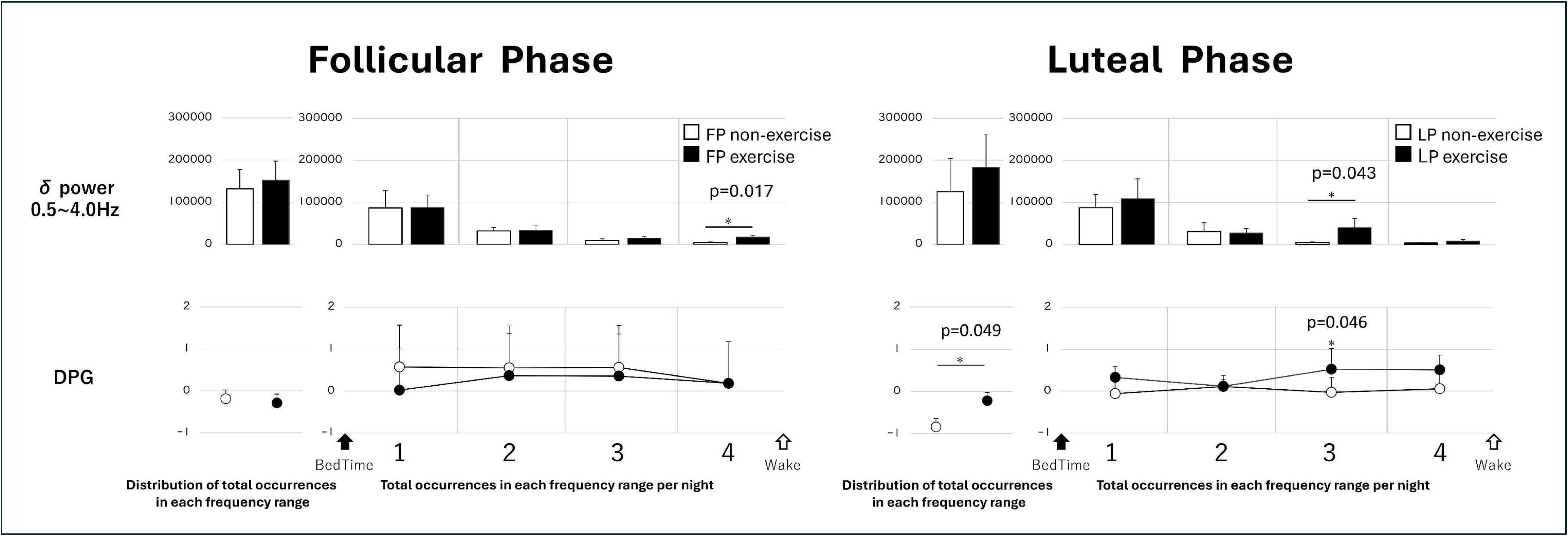
Effects of exercise on EEG spectral power and DPG during sleep across menstrual phases. Cumulative δ power (0.5–4.0 Hz) and distal–proximal skin temperature gradient (DPG) during nocturnal sleep in the follicular (left panels) and luteal (right panels) phases under non-exercise (open bars) and exercise (filled bars) conditions. Each left bar represents the total δ power or mean DPG across the entire sleep period, while the right grouped bars indicate the distribution or progression across sleep segments (e.g., early, middle, late night), reflecting temporal patterns throughout the night. Values are expressed as mean ± SE. Asterisks and p-values denote statistically significant differences between conditions (\**p <* 0.05).

### Subjective sleep parameters upon waking

Table 3 shows the subjective sleep indices assessed upon waking. In the luteal phase, the items “difficulty in concentrating” and “tired eyes” were significantly improved in the exercise condition compared with the non-exercise condition (z = 2.060, *p* = 0.039 and z = 2.041, *p* = 0.041, respectively). No significant differences in the other subjective sleep parameters were observed.

**Table 3.**
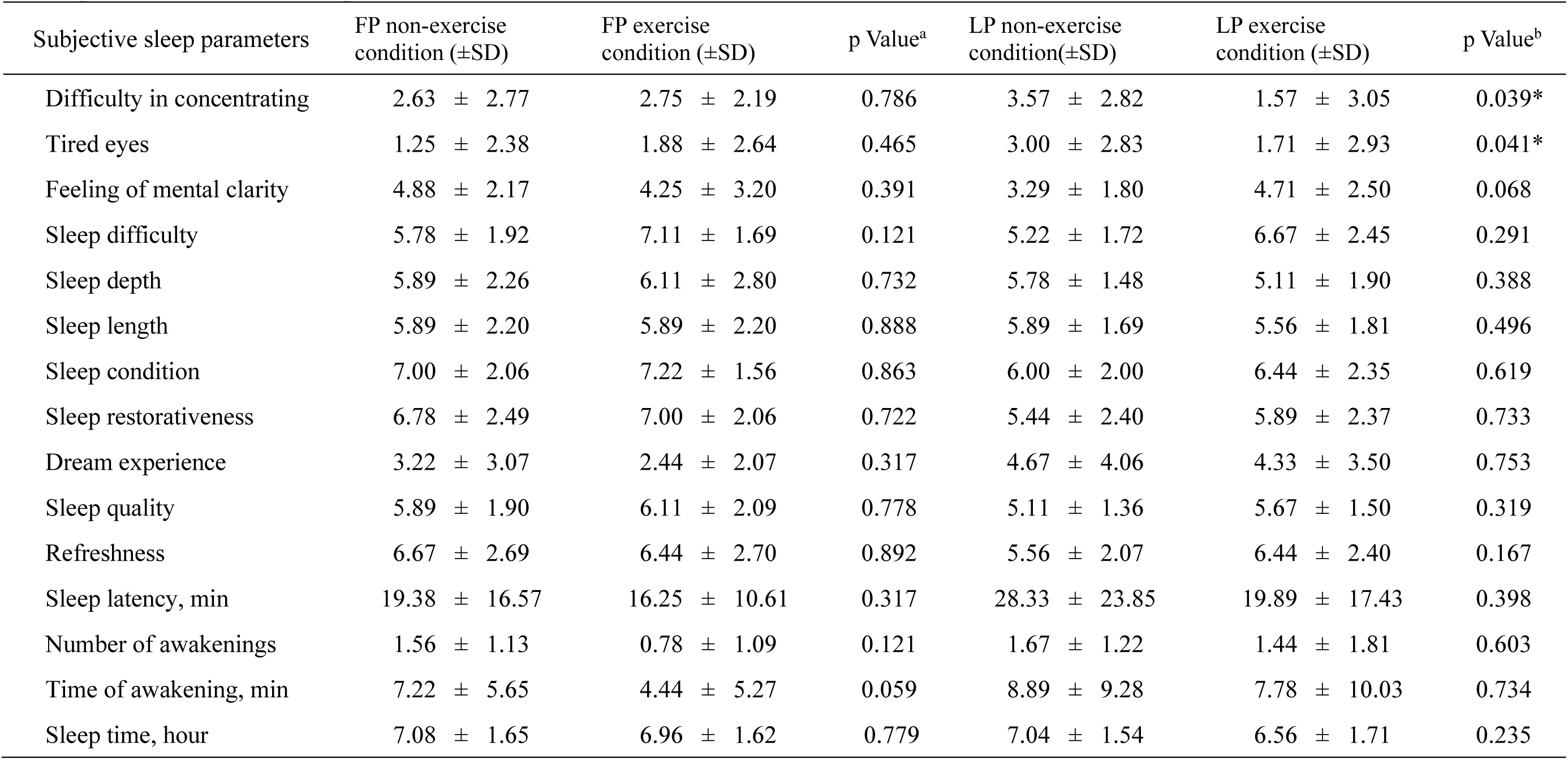
Subjective sleep parameters under non-exercise and exercise conditions during the follicular and luteal phases. Mean values (± SD) of subjective sleep-related variables assessed under non-exercise and exercise conditions during the follicular and luteal phases(*p <* 0.05). a Comparison between follicular-phase non-exercise and exercise conditions. b Comparison between luteal-phase non-exercise and exercise conditions.

| Subjective sleep parameters | FP non-exercise<br>condition (±SD) | FP exercise<br>condition (±SD) | p Value<sup>a</sup> | LP non-exercise<br>condition(±SD) | LP exercise<br>condition (±SD) | p Value<sup>b</sup> |
| --- | --- | --- | --- | --- | --- | --- |
| Difficulty in concentrating | 2.63 ± 2.77 | 2.75 ± 2.19 | 0.786 | 3.57 ± 2.82 | 1.57 ± 3.05 | 0.039* |
| Tired eyes | 1.25 ± 2.38 | 1.88 ± 2.64 | 0.465 | 3.00 ± 2.83 | 1.71 ± 2.93 | 0.041* |
| Feeling of mental clarity | 4.88 ± 2.17 | 4.25 ± 3.20 | 0.391 | 3.29 ± 1.80 | 4.71 ± 2.50 | 0.068 |
| Sleep difficulty | 5.78 ± 1.92 | 7.11 ± 1.69 | 0.121 | 5.22 ± 1.72 | 6.67 ± 2.45 | 0.291 |
| Sleep depth | 5.89 ± 2.26 | 6.11 ± 2.80 | 0.732 | 5.78 ± 1.48 | 5.11 ± 1.90 | 0.388 |
| Sleep length | 5.89 ± 2.20 | 5.89 ± 2.20 | 0.888 | 5.89 ± 1.69 | 5.56 ± 1.81 | 0.496 |
| Sleep condition | 7.00 ± 2.06 | 7.22 ± 1.56 | 0.863 | 6.00 ± 2.00 | 6.44 ± 2.35 | 0.619 |
| Sleep restorativeness | 6.78 ± 2.49 | 7.00 ± 2.06 | 0.722 | 5.44 ± 2.40 | 5.89 ± 2.37 | 0.733 |
| Dream experience | 3.22 ± 3.07 | 2.44 ± 2.07 | 0.317 | 4.67 ± 4.06 | 4.33 ± 3.50 | 0.753 |
| Sleep quality | 5.89 ± 1.90 | 6.11 ± 2.09 | 0.778 | 5.11 ± 1.36 | 5.67 ± 1.50 | 0.319 |
| Refreshness | 6.67 ± 2.69 | 6.44 ± 2.70 | 0.892 | 5.56 ± 2.07 | 6.44 ± 2.40 | 0.167 |
| Sleep latency, min | 19.38 ± 16.57 | 16.25 ± 10.61 | 0.317 | 28.33 ± 23.85 | 19.89 ± 17.43 | 0.398 |
| Number of awakenings | 1.56 ± 1.13 | 0.78 ± 1.09 | 0.121 | 1.67 ± 1.22 | 1.44 ± 1.81 | 0.603 |
| Time of awakening, min | 7.22 ± 5.65 | 4.44 ± 5.27 | 0.059 | 8.89 ± 9.28 | 7.78 ± 10.03 | 0.734 |
| Sleep time, hour | 7.08 ± 1.65 | 6.96 ± 1.62 | 0.779 | 7.04 ± 1.54 | 6.56 ± 1.71 | 0.235 |

### Correlation between heat dissipation and subjective sleep quality

Figure 5 shows the correlation between physiological and subjective sleep parameters. In the luteal phase under the exercise condition, a significant negative correlation was found between pre-sleep heat dissipation (indexed by the DPG) and physical fatigue before bedtime (*p* = 0.020, Spearman’s ρ = −0.883). This finding suggested that individuals with greater thermoregulatory heat loss experienced lower levels of physical fatigue. Additionally, in the luteal phase, a significant negative correlation was observed between the percentage of stage N3 sleep and difficulty in concentrating upon waking (*p* = 0.030, ρ = −0.802). This finding indicated that greater deep sleep was associated with improved cognitive clarity upon awakening.

**Figure 5.**
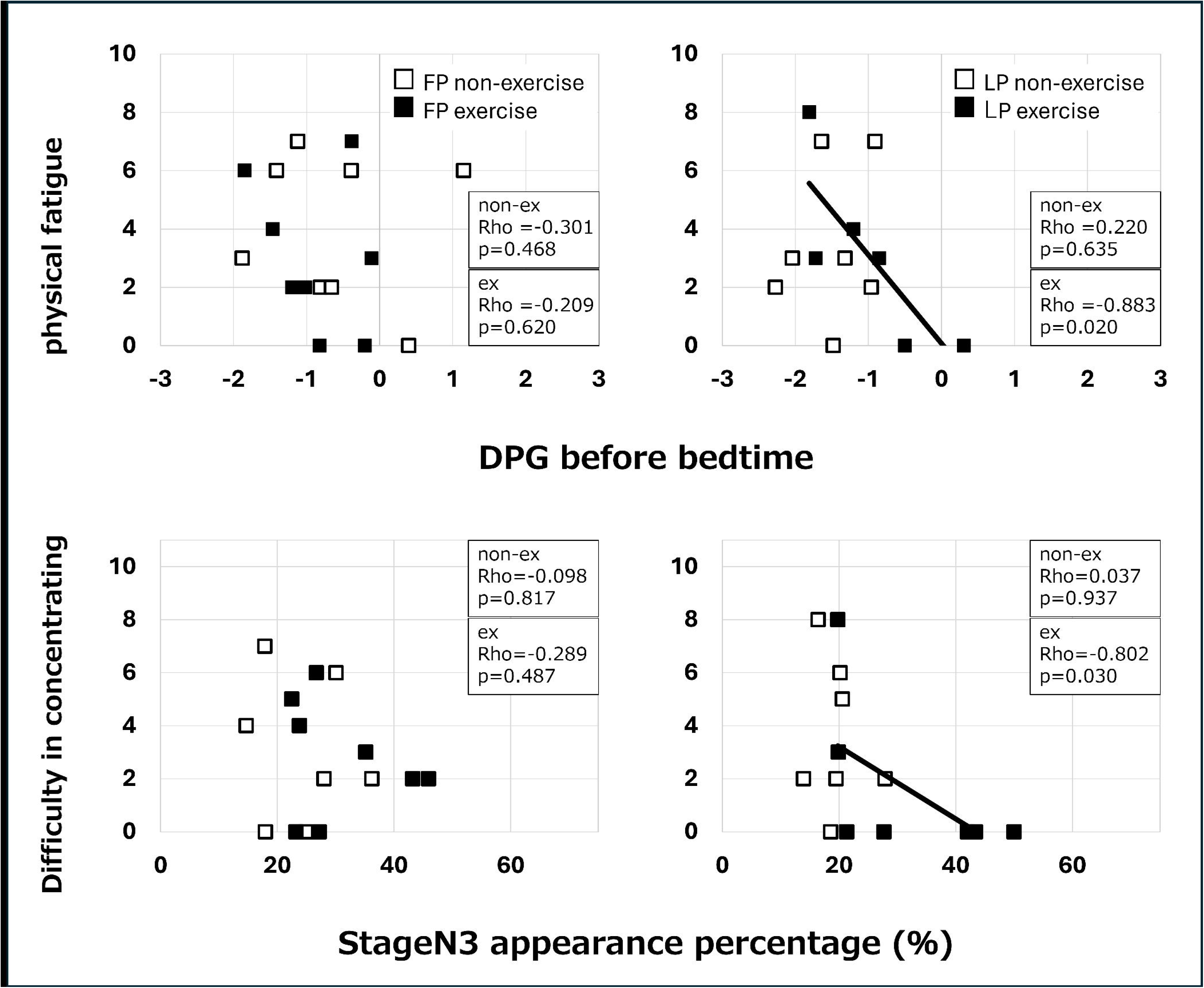
Associations between thermoregulatory/sleep parameters and subjective symptoms in each menstrual phase. Scatterplots showing correlations between distal–proximal skin temperature gradient (DPG) before bedtime and subjective physical fatigue (top row), and between stage N3 sleep percentage and difficulty in concentrating (bottom row), under non-exercise (open squares) and exercise (filled squares) conditions. Left panels represent data from the follicular phase; right panels represent data from the luteal phase. Spearman’s rank correlation coefficients (Rho) and p-values are shown for each condition. Solid and dashed lines indicate trend lines for the exercise and non-exercise conditions, respectively.

## DISCUSSION

This study is, to our knowledge, the first to demonstrate that a single session of moderate-intensity resistance training enhances nocturnal heat dissipation and deep sleep particularly in the luteal phase, when thermoregulatory function is typically impaired. Previous studies have shown that habitual exercisers have enhanced heat dissipation responses, especially in the luteal phase[12]. Our previous study also showed that daytime exercise facilitated thermoregulatory heat loss and increased deep sleep in healthy young men[11]. On the basis of these findings, we chose resistance training as an accessible, non-equipment-based intervention to improve sleep by enhanced heat dissipation. The time of exercise (14:00–14:30 hours) was selected on the basis of previous reports suggesting that physical activity 2–4 hours before bedtime does not disrupt sleep[23] and that resistance training timing does not affect the total sleep time or REM duration. A 70% one-repetition maximum intensity was adopted in accordance with reports showing that this level can improve objective sleep indices. The 30-minute full-body protocol was verified for feasibility in young women without exercise habits under the guidance of an athletic trainer.

In this study, in both menstrual phases, resistance training increased nocturnal deep sleep. However, in the luteal phase, the DPG remained elevated from pre-sleep through the night, and the proportion of stage N3 sleep remained high in the latter half of sleep. These results are consistent with our previous study, which showed that an increased DPG was associated with enhanced slow-wave sleep. Progesterone elevates the core body temperature[2] and enhances skin vasodilation sensitivity[24]. Therefore, exercise in the luteal phase may facilitate thermoregulatory heat loss and subsequently deepen sleep. This possibility is supported by findings from a study by Ogura et al. who reported enhanced thermoregulatory responses to exercise in the luteal phase[12]. The increase in δ power observed along with an elevated DPG suggests a reciprocal relationship between thermoregulation and deep sleep. According to McGinty and Szymusiak, slow-wave sleep may contribute to heat loss by decreasing metabolic heat production and promoting vasodilation and sweating[25]. In this context, our findings suggest that not only does exercise facilitate pre-sleep heat loss but deep sleep itself may further aid thermoregulation during the night. Although our resistance training protocol involved full-body movement, the participants reported higher perceived exertion during upper body exercises, such as push-ups. This exertion may have contributed to increased heat dissipation from the hands. A previous study showed that leg-based ergometer exercise increased the DPG from the feet [11], suggesting that regional differences in thermoregulatory responses depend on the muscles engaged. However, further studies are required to clarify the relationship between exercise modality and regional heat dissipation.

In addition to improvements in objective sleep parameters, our study showed that physiological markers, particularly the percentage of stage N3 sleep and pre-sleep DPG, were significantly associated with subjective sleep quality upon waking. Specifically, in the luteal phase exercise condition, a higher proportion of stage N3 sleep was correlated with reduced difficulty in concentrating upon waking, and an increased pre-sleep DPG was associated with lower levels of physical fatigue at bedtime. Although group-level differences in most subjective sleep indices were not significant, these correlations suggested that individuals who experienced greater thermoregulatory responses and deeper sleep also perceived better mental and physical recovery. This finding indicates the potential of resistance training not only to alter sleep architecture but also to improve subjective well-being in a physiologically meaningful way.

This study has several limitations. First, hormonal measurements were not conducted to confirm menstrual phases. The basal body temperature and ovulation kits were used for phase identification instead. Second, nocturnal EEG was recorded using a portable device in participants’ homes rather than in a supervised sleep laboratory setting, which limited our ability to control for environmental factors or respond to electrode disconnections. Third, tympanic membrane temperature was used as a proxy for core body temperature, although it is more sensitive to environmental fluctuations than rectal measurements and may introduce variability. Future studies should include hormonal verification of menstrual phases, conduct laboratory-based sleep monitoring, and examine chronic resistance training interventions. Additionally, comparing the effects of various exercise types, intensities, and timing on objective and subjective sleep quality is essential to clarify optimal strategies for improving sleep in women.

In conclusion, moderate intensity resistance training during the day enhances thermoregulatory heat loss and increases deep sleep, especially in the luteal phase when heat dissipation is typically impaired. These findings suggest that body weight resistance training may serve as a simple and effective intervention to promote sleep quality in young women across different menstrual phases.

## ACKNOWLEDGMENTS

We thank the members of our research laboratory, namely Mizuki Kiyama, Himeka Kubokawa, Konomi Sugawara, and Mariko Takakura, for their cooperation in the experiments and helpful discussions, which contributed greatly to the completion of this project. We would also like to thank the individuals who participated in this study. We thank Ellen Knapp, PhD, from Edanz (https://jp.edanz.com/ac) for editing a draft of this manuscript.

## AUTHORS’ CONTRIBUTIONS

S.A.-O., M.F., and S.N. conceived and designed the study; M.F., R.I., and S.A.-O. performed the experiments; M.F., R.I., and S.A.-O. analyzed the data; M.F. and S.A.-O. interpreted the results; M.F. and S.A.-O. wrote the manuscript; S.A.-O. obtained funding; S.A.-O. supervised; M.F., R.I., S.N., A.K., K.K., M.T. and S.A.-O. reviewed and approved the final version of the manuscript.

## References

1. Shechter A, Boivin DB. Sleep, Hormones, and Circadian Rhythms throughout the Menstrual Cycle in Healthy Women and Women with Premenstrual Dysphoric Disorder. Int J Endocrinol. 2010;2010: 259345.

2. Baker FC, Siboza F, Fuller A. Temperature regulation in women: Effects of the menstrual cycle. Temperature (Austin). 2020;7: 226–262.

3. Shibui K, Uchiyama M, Okawa M, Kudo Y, Kim K, Liu X, et al. Diurnal fluctuation of sleep propensity and hormonal secretion across the menstrual cycle. Biol Psychiatry. 2000;48: 1062–1068.

4. Manber R, Bootzin RR. Sleep and the menstrual cycle. Health Psychol. 1997;16: 209–214.

5. Sharkey KM, Crawford SL, Kim S, Joffe H. Objective sleep interruption and reproductive hormone dynamics in the menstrual cycle. Sleep Med. 2014;15: 688– 693.

6. Baker FC, Driver HS. Circadian rhythms, sleep, and the menstrual cycle. Sleep Med. 2007;8: 613–622.

7. Heinemann LAJ, Do Minh T, Filonenko A, Uhl-Hochgräber K. Explorative evaluation of the impact of premenstrual disorder on daily functioning and quality of life. Patient. 2010;3: 125–132.

8. Baker FC, Lamarche LJ, Iacovides S, Colrain IM. Sleep and menstrual-related disorders. Sleep Med Clin. 2008;3: 25–35.

9. Erbil N, Yücesoy H. Relationship between premenstrual syndrome and sleep quality among nursing and medical students. Perspect Psychiatr Care. 2022;58: 448–455.

10. Kovacevic A, Mavros Y, Heisz JJ, Fiatarone Singh MA. The effect of resistance exercise on sleep: A systematic review of randomized controlled trials. Sleep Med Rev. 2018;39: 52–68.

11. Aritake-Okada S, Tanabe K, Mochizuki Y, Ochiai R, Hibi M, Kozuma K, et al. Diurnal repeated exercise promotes slow-wave activity and fast-sigma power during sleep with increase in body temperature: a human crossover trial. J Appl Physiol. 2019;127: 168–177.

12. Yukio Ogura, Tomoko Kuwahara, Yoshimitsu Inoue. Effects of Physical Training on Heat Loss Responses in Young Women. DESCENTE SPORTS SCIENCE. 2003. pp. 86–95.

13. Kuwahara T, Inoue Y, Taniguchi M, Ogura Y, Ueda H, Kondo N. Effects of physical training on heat loss responses of young women to passive heating in relation to menstrual cycle. Eur J Appl Physiol. 2005;94: 376–385.

14. Maged AM, Abbassy AH, Sakr HRS, Elsawah H, Wagih H, Ogila AI, et al. Effect of swimming exercise on premenstrual syndrome. Arch Gynecol Obstet. 2018;297: 951–959.

15. Buysse DJ, Reynolds CF 3rd, Monk TH, Berman SR, Kupfer DJ. The Pittsburgh Sleep Quality Index: a new instrument for psychiatric practice and research. Psychiatry Res. 1989;28: 193–213.

16. Goldberg DP, Blackwell B. Psychiatric illness in general practice. A detailed study using a new method of case identification. Br Med J. 1970;1: 439–443.

17. Radloff LS. The CES-D Scale: A Self-Report Depression Scale for Research in the General Population. Appl Psychol Meas. 1977;1: 385–401.

18. Moos RH. The development of a menstrual distress questionnaire. Psychosom Med. 1968;30: 853–867.

19. Lee CK, Lee J-H, Ha M-S. Comparison of the Effects of Aerobic versus Resistance Exercise on the Autonomic Nervous System in Middle-Aged Women: A Randomized Controlled Study. Int J Environ Res Public Health. 2022;19. doi:10.3390/ijerph19159156

20. Berry RB, Albertario CL, Harding SM, Lloyd RM, Plante DT, Quan SF, et al. The AASM manual for the scoring of sleep and associated events: rules, terminology and technical specifications. American Academy of Sleep Medicine; 2018.

21. Kräuchi K, Cajochen C, Werth E, Wirz-Justice A. Functional link between distal vasodilation and sleep-onset latency? Am J Physiol Regul Integr Comp Physiol. 2000;278: R741–8.

22. Hoddes E, Zarcone V, Smythe H, Phillips R, Dement WC. Quantification of sleepiness: a new approach. Psychophysiology. 1973;10: 431–436.

23. Frimpong E, Mograss M, Zvionow T, Dang-Vu TT. The effects of evening high-intensity exercise on sleep in healthy adults: A systematic review and meta-analysis. Sleep Med Rev. 2021;60: 101535.

24. Inoue Y, Tanaka Y, Omori K, Kuwahara T, Ogura Y, Ueda H. Sex– and menstrual cycle-related differences in sweating and cutaneous blood flow in response to passive heat exposure. Eur J Appl Physiol. 2005;94: 323–332.

25. McGinty D, Szymusiak R. Keeping cool: a hypothesis about the mechanisms and functions of slow-wave sleep. Trends Neurosci. 1990;13: 480–487.

